# BioDiscViz : a visualization support and consensus signature selector for BioDiscML results

**DOI:** 10.1101/2022.10.07.511250

**Authors:** Sophiane Bouirdene, Mickael Leclercq, Léopold Quitté, Steve Bilodeau, Arnaud Droit

**Affiliations:** Département de Médecine Moléculaire CHU de Québec Research Center, Université Laval, Québec, QC, Canada; Département de biologie moléculaire, biochimie médicale et pathologie, Faculté de Médecine, Université Laval, Québec, Canada

## Abstract

Machine learning (ML) algorithms are powerful tools to find complex patterns and biomarker signatures where conventional statistical methods may fail to identify them. While the ML field made significant progress, state of the art methodologies to build efficient and non-overfitting models are not always applied in the litterature. To this purpose, automatic programs, such as BioDiscML, have been designed to identify biomarker signatures and correlated features while escaping overfitting using multiple evaluation strategies, such as cross validation, bootstrapping and repeated holdout. To further improve BioDiscML and reach a broader audience, better visualization support and flexibility in choosing the best models and signatures are needed. Thus, to provide researchers with an easily accessible and usable tool for in depth investigation of the results from BioDiscML outputs, we developed a visual interaction tool called BioDiscViz. This tool provides summaries, tables and graphics, in the form of Principal Component Analysis (PCA) plots, heatmaps and boxplots for the best model and the correlated features. Furthermore, this tool also provides visual support to extract a consensus signature from BioDiscML models using a combination of filters. BioDiscViz will be a great visual support for research implying machine learning, hence new opportunities in this field by opening it to a broader community.

## Introduction

In recent years, new methods of Artificial Intelligence (AI) have been deployed in bioinformatics research to provide pattern classification, biomarker identification and forecast modeling using omics data. Studying biomarker signatures is an important part of the research process as they are correlated to biological functions. Machine learning and feature selection will identify multivariate associations of biomarkers (i.e., features) and detect complex hidden patterns in the data. Considering the existence of many algorithms for feature selection and classification, multiple models are often generated with different signatures, but inconsistent overlaps between signatures despite equivalent performances are frequent (Li et al. 2016). Furthermore, correlated features may not be retained by the models during their optimization when avoiding redundancy of information. Indeed, selecting a “best model” and its signatures is an equilibrium between decomplexifying the model and getting all valuable biomarkers. Often, various approaches, like ensemble learning or union of overlapping features, tend to find optimized solutions but at the cost of either side of the balance.

A solution to facilitate the generation of multiple models and signatures has been proposed with an automatic ML tool, BioDiscML (Leclercq et al. 2019). BioDiscML is a new generation ML tool which has been demonstrated to be highly efficient in multiple research topics involving the identification of biomarker signatures from various types of data, such as proteomics (Roux-Dalvai et al. 2019), transcriptomics (Rabaglino and Kadarmideen 2020) and multi-omics (metagenomics/metabolomics, metagenomics/lipidomics) (Khorraminezhad et al. 2020; Doré et al. 2022). Furthermore, BioDiscML proposes various conditions for choosing a “best model” from its results, but this is complex to determine as some data are too heterogeneous to propose ideal decision threshold metrics. However, this tool does not provide visualization of the signature, hence limiting a rapid view of the results. Thus, to help in these decisions, we propose a visual tool, BioDiscViz, to support the choice of consensus features within a set of trained classifiers with their corresponding signatures.

## Design and implementation

BioDiscViz is a visual Shiny application working on Windows and Unix operating systems to support BioDiscML by presenting an interactive interface and graphs to the researchers which will improve their understanding of the results. The application is based on R (R Core Team 2021). It uses the framework Rshiny (Chang W et al 2021) and its dependency Rshiny Dashboard (Chang W, Borges Ribeiro B 2021) and requires Rstudio (RStudio Team 2020), an integrated development environment for R.

### Input

BioDiscViz takes as input a directory containing BioDiscML output in csv format and their summary results. The best model and the classification or regression results are independently accessible. Furthermore, the tool supports multiple BioDiscML outputs in the same directory and allows rapid switching between them.

### Layout

The application is divided into four main parts which are : short signature, long signature, attribute distribution and consensus signature.

*Short signature* presents the results of the best model selected by BioDiscML in the form of different plots representing the features selected by the models.

*Long signature* gives a representation of the correlated biomarkers.

*Attribute distribution* is an addition of the visualizer which allows to interactively visualize the features that were used the most by the different classifiers tested. The user is able to select different thresholds for the classifiers using the Matthew correlation coefficient and standard deviation metrics. Furthermore, the user determines the number of attributes to match the experimental design.

*Consensus signature* gives a representation of the different signatures which were called by the most classifiers under user-determined parameters.

### Representations

There is a heatmap, a PCA graph and a boxplot to represent the short, long and consensus signatures. The heatmap was made using ComplexHeatmap (Gu, Eils, and Schlesner 2016), an user-friendly package for better representation of heatmaps. The PCA and boxplots were built using FactoExtra (Kassambara 2020) and ggplot2 (Wickham H 2016) respectively.

The attribute distribution is represented under the form of a UpsetR plot (Conway, Lex, and Gehlenborg 2017) (Fig1.B). UpsetR is a R package generating static upset plots to visualize the intersections between the different features in the different classifiers.

**Figure 1:**
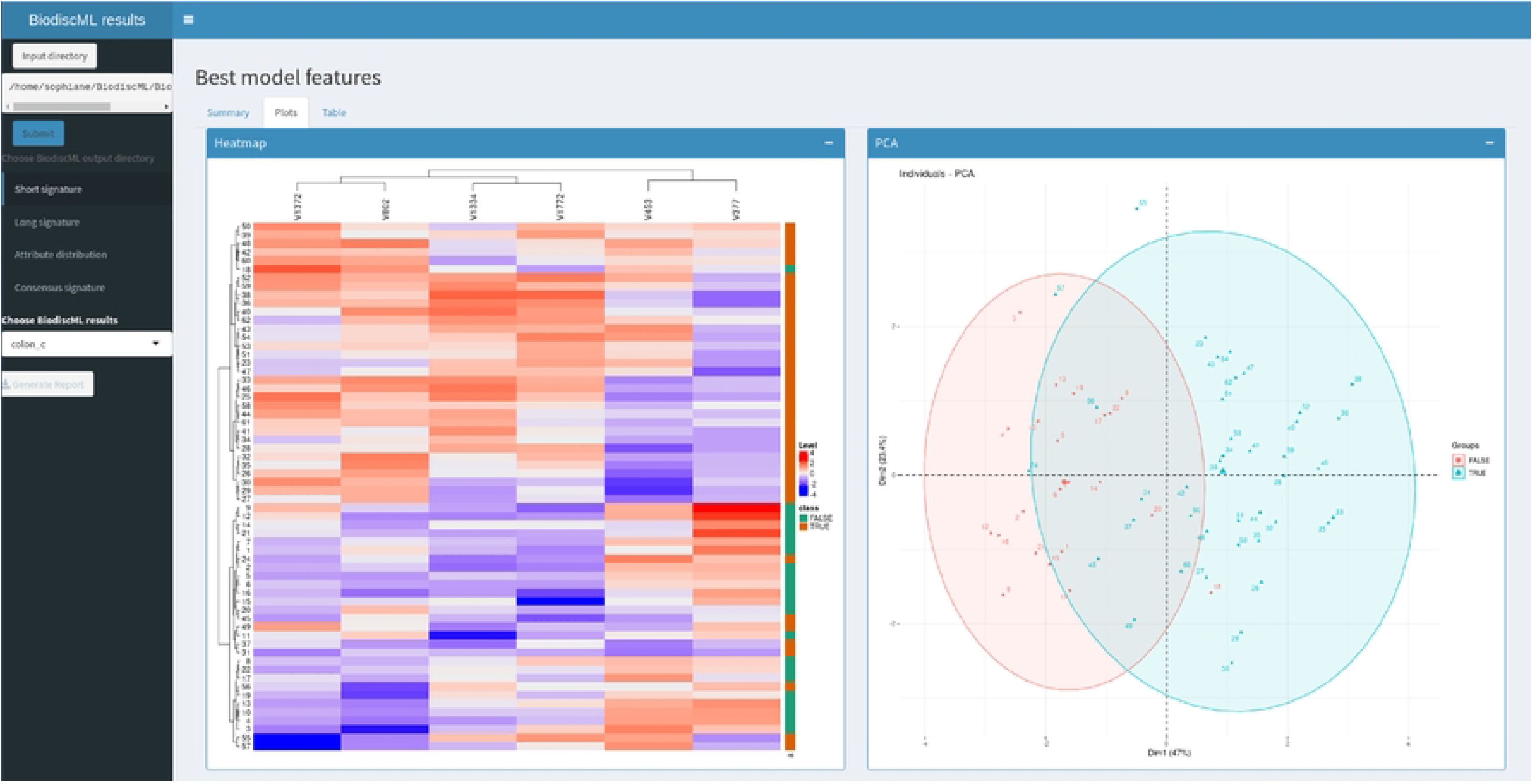
**A.** Heatmap and PCA by BiodiscViz on the best model found by BiodiscML, Kstar model,on the colon cancer dataset. We observe in cyan color the tumor tissues and in red the normal ones. **B**. Selection of the consensus signatures with BiodiscViz on the colon cancer dataset. Here were selected the 10 attributes most frequently called by the classifiers passing the threshold of a Matthew Correlation Coefficient >= 0.75 and a Standard Deviation <= 0.15.

BioDiscViz also gives access to the summary details for the short signature and an interactive table of the data used for the short and long signature.

Considering that non-numerical features cannot be easily integrated into PCA and heatmap with other numerical values, a particularity of BioDiscViz is the transformation of categorical features into numerical ones. This form allows one to simply annotate them on the side of the heatmaps to integrate the information contained by these features into the clustering of PCA.

### Outputs

BioDiscViz also possesses different functions to facilitate use and export of the results for archiving, sharing and publication. The first one is the creation of a report of the different graphs represented in the application and takes into account the modifications carried out by the user. The second functionality is to be able to download a sub dataset containing the information for the selected features in the “attribute distribution”.

The study of consensus signatures has a great interest as it allows researchers to identify new molecular targets of interest. If the best model provides a vision of which useful data were selected, the model does not necessarily use the features providing the most information. As we consider that the most frequently used signatures by the classifiers contain important information for our problem. Forcing BioDiscML’s models to use these consensus signatures could potentially improve the initial results obtained by the previous best model.

## Results

To demonstrate the functionalities of BioDiscViz, we used a colon cancer dataset (Alon et al. 1999) which was used for the BioDiscML publication and which is available on BioDiscViz gitlab. This dataset contains gene expression in 40 tumor and 22 normal colon tissue samples.

### Visualize the best signature

The identification of the best signatures were studied from two perspectives. First, the signatures from the best model followed by the consensus signatures.

For signatures retrieved from the best model, different plots were generated. In this case, two classes were cleanly separated on the PCA and the heatmap (Fig 1.A), showing that they provide enough information to the model to correctly predict tumor tissues and healthy tissues. Then, differential expressions of genes identified in the model were visualized using boxplots (Fig.2). Interestingly, all the signatures showed promising results as there is a clear difference for each gene between the two classes.

**Figure 2:**
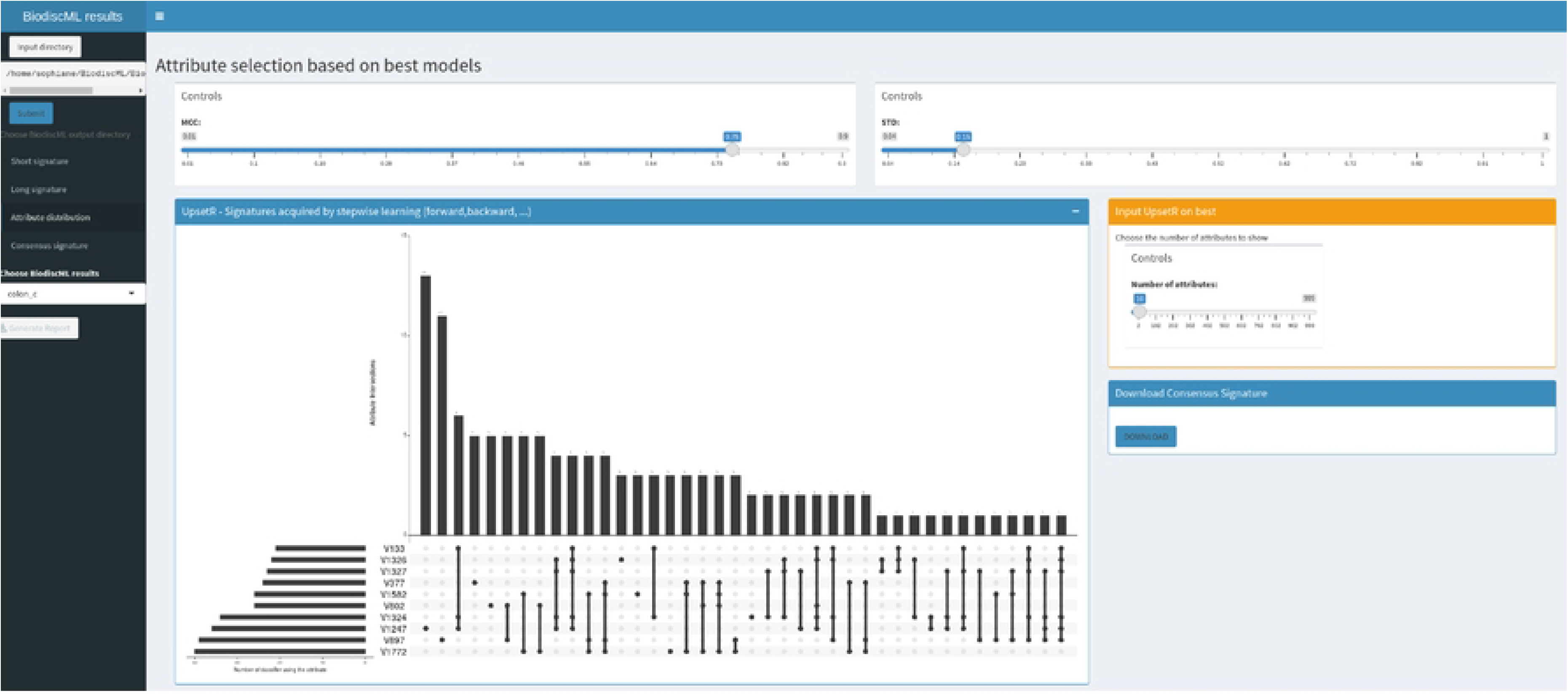
Boxplot of the best model obtained by BioDiscML

### Consensus signature

BioDiscML uses ensemble methods to create an association of signatures, but it does not take advantage of all generated models during its learning stage. Ensemble methods also keep all features of the model’s signatures, without any optimization, thus complexifying the model. We made the hypothesis that the features which were called the most frequently by the different models were containing useful information for the problem treated.These signatures, which we call consensus signatures, could represent an interesting option to rebuild a new signature and improve or simplify the model. To look further into the signatures, it is possible to generate a dataset of consensus signatures directly with BioDiscViz. After different tests, it was decided to focus on the 10 best signatures obtained from the classifiers which were passing the threshold of MCC>=0.76 (Matthew’s Correlation Coefficient) and STD_MCC<=0.15 (standard deviation of MCC) (Fig.1B). The quality of the selection was assessed using the heatmap (Fig.3A) and PCA (Fig. 3B), which presented a better separation between the classes than the best model identified by BioDiscML. Compared to the best model signatures, these consensus signatures consist of 3 genes overlapping with the best signature, and 7 genes have been newly added. To further look into these new signatures it was possible to use the boxplot to only select genes which seemed to have a clear separation in the distributions between the tissues with and without cancer.

**Figure 3:**
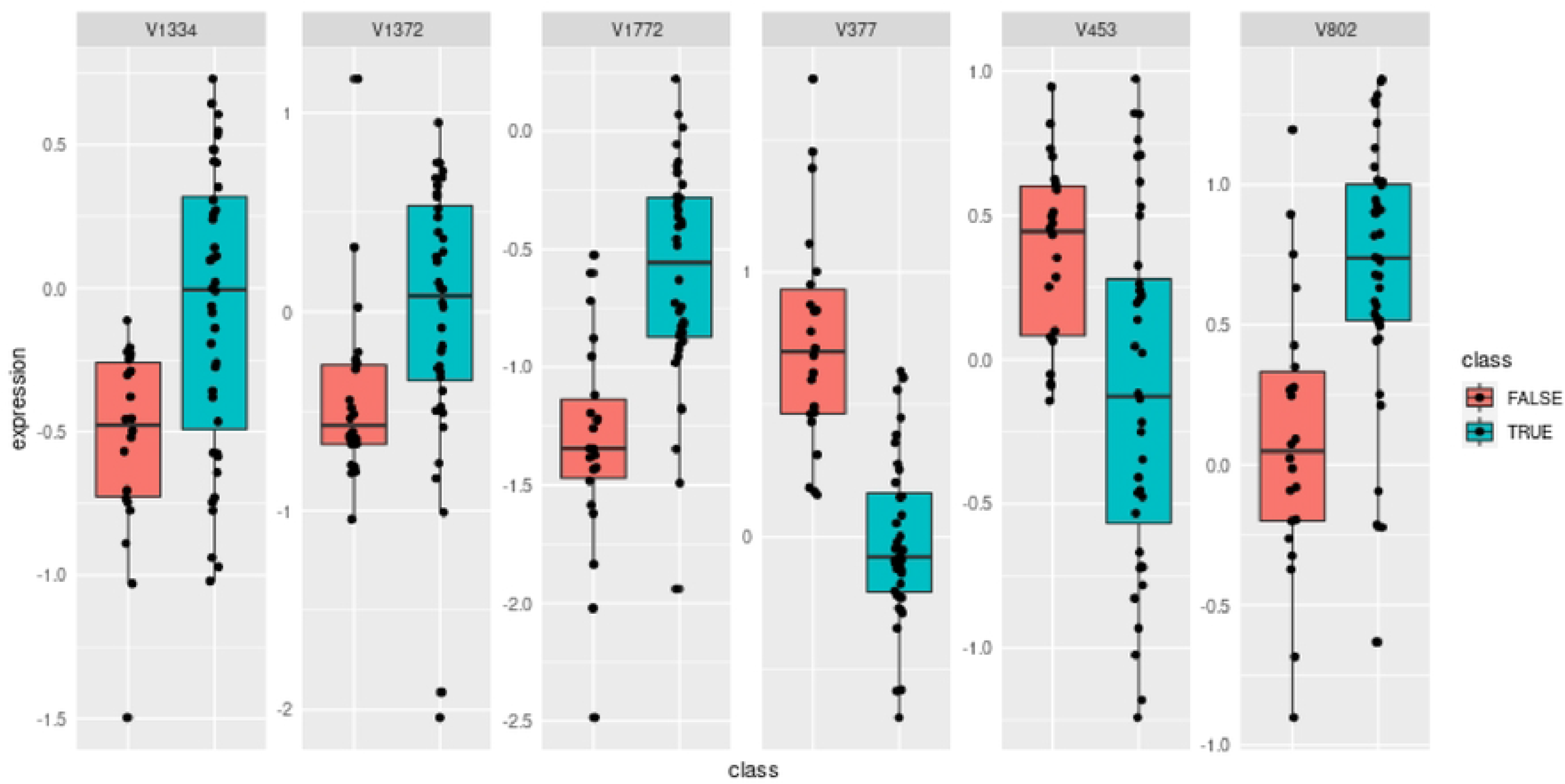
Graphical representation of the 10 best consensus signatures **A**. Heatmap. **B**. PCA

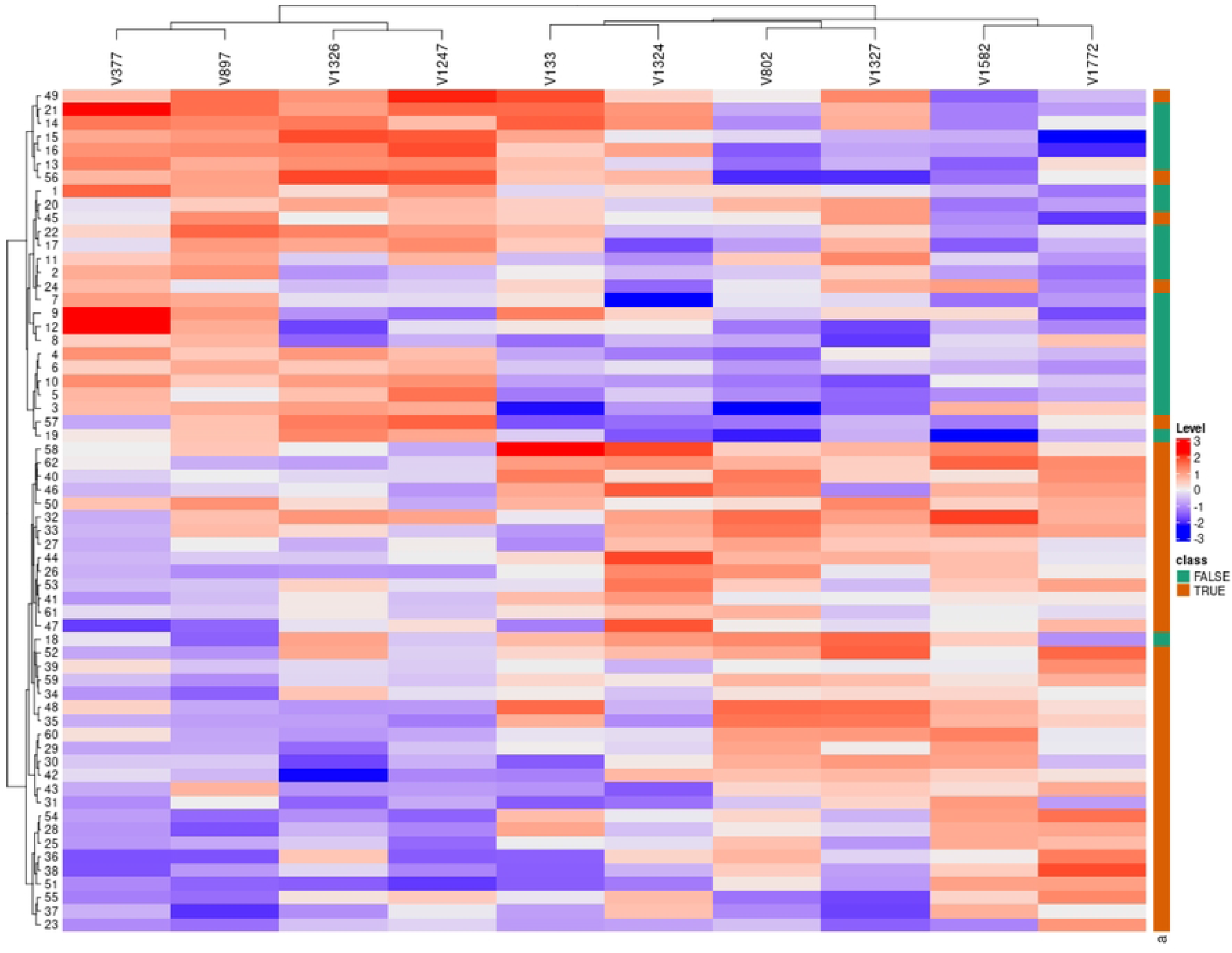

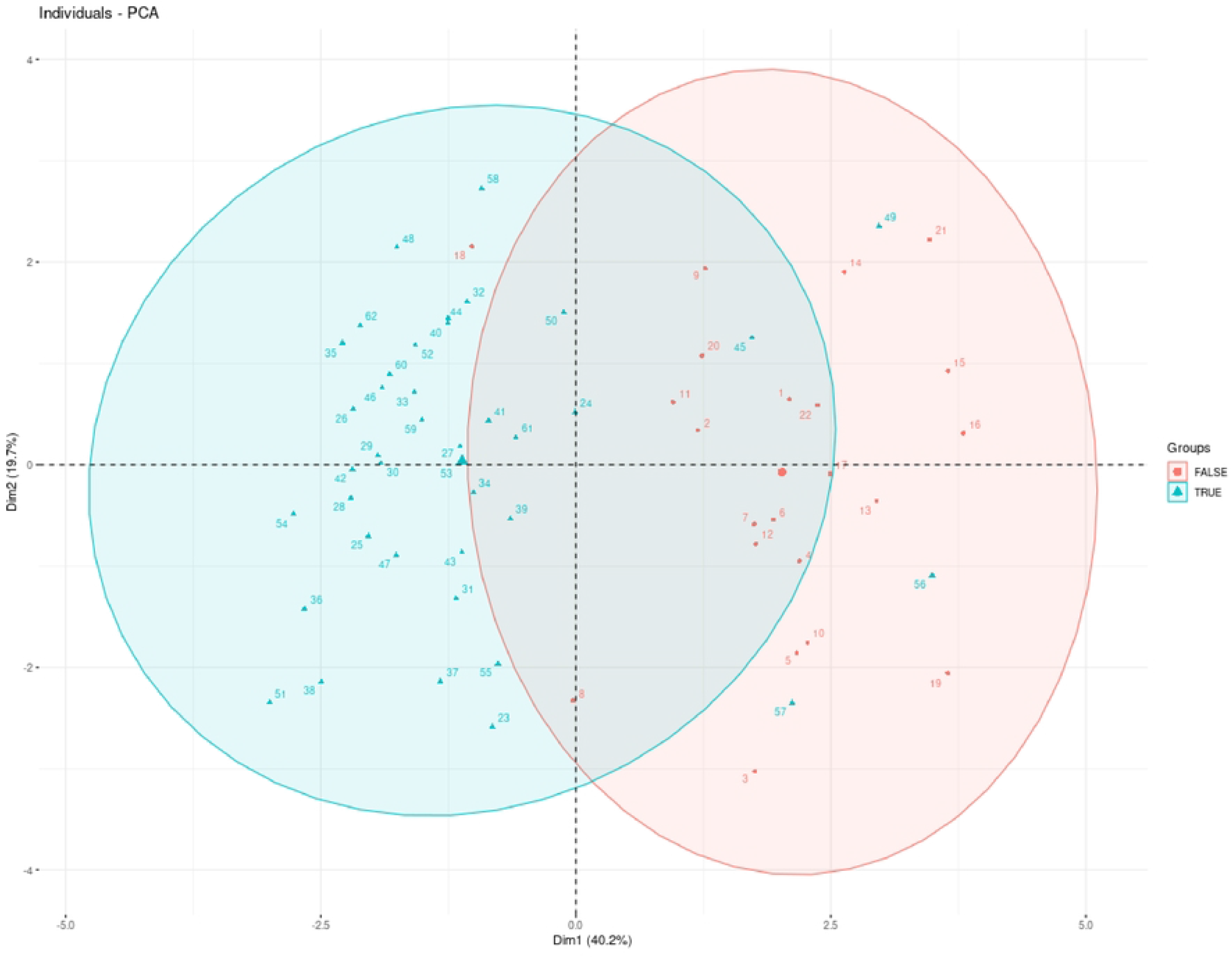

Following the identification of the consensus signature, we re-runned BioDiscML to find an optimal machine learning classifier with the full signature, without any feature selection. The best classifier was a Kstar model with 6 attributes signature which had a MCC of 0.776 with a standard deviation across (STD) all evaluation procedures of 0.037, which is a reasonable performance considering past work on MCC evaluations (Schober, Boer, and Schwarte 2018). Furthermore, the model had an accuracy of 0.8571 which is comparable or even better compared to the literature for this type of data (“Detection of Effective Genes in Colon

Cancer: A Machine Learning Approach” 2021). With the consensus signature, obtained a Fuzzy Lattice Reasoning model with a MCC of 0.791 (STD 0.032) which is slightly better than the previous best model (MCC increased by 1,9% and STD decreased by 15,6%).

In Conclusion our tool is able to give visual support to BioDiscML and new insights outside of the best model by looking into the consensus signatures. Furthermore, these consensus signatures could be used to rerun BioDiscML and may enhance the quality of the model.

## Availability and Future Directions

Biodisc-Viz is directly implemented in R and can be downloaded from Gitlab (https://gitlab.com/SBouirdene/biodiscviz.git)

